# Evolutionary genetics of cytoplasmic incompatibility genes *cifA* and *cifB* in prophage WO of *Wolbachia*

**DOI:** 10.1101/180075

**Authors:** Amelia R. I. Lindsey, Danny W. Rice, Sarah R. Bordenstein, Andrew W. Brooks, Seth R. Bordenstein, Irene L. G. Newton

## Abstract

The bacterial endosymbiont *Wolbachia* manipulates arthropod reproduction to facilitate its maternal spread through populations. The most common manipulation is cytoplasmic incompatibility (CI): *Wolbachia*-infected males produce modified sperm that cause embryonic mortality, unless rescued by eggs harboring the same *Wolbachia*. The genes underlying CI, *cifA* and *cifB,* were recently identified in the eukaryotic association module of *Wolbachia*’s prophage WO. Here, we use transcriptomic and genomic approaches to address three important evolutionary facets of these genes. First, we assess whether or not *cifA* and *cifB* comprise a classic toxin-antitoxin operon, and show they do not form an operon in strain *w*Mel. They coevolve but exhibit strikingly distinct expression across host development. Second, we provide new domain and functional predictions across homologs within *Wolbachia*, and we show amino acid sequences vary substantially across the genus. Lastly, we investigate conservation of *cifA* and *cifB* and find degradation and loss of the genes is common in strains that no longer induce CI. Taken together, we find no evidence for the operon hypothesis in *w*Mel, provide functional annotations that broaden the potential mechanisms of CI induction, illuminate recurrent erosion of *cifA* and *cifB* in non-CI strains, and advance an understanding of the most widespread form of reproductive parasitism.

## Introduction

The genus *Wolbachia* is the most widespread group of maternally transmitted endosymbiotic bacteria (Zug and Hammerstein 2012). They occur worldwide in numerous arthropods and nematodes and can selfishly manipulate reproduction (Werren, et al. 2008), confer antiviral defense (Bian, et al. 2010; Teixeira, et al. 2008), and assist reproduction and development of their hosts (Dedeine, et al. 2001; Hoerauf, et al. 1999; Hosokawa, et al. 2010). The most common parasitic manipulation is cytoplasmic incompatibility (CI), whereby *Wolbachia*-infected males produce modified sperm that can only be rescued by eggs infected with the same *Wolbachia* strain (Yen and Barr 1971). If the modified sperm fertilize eggs infected with no *Wolbachia* (unidirectional CI) or a genetically-incompatible *Wolbachia* strain (bidirectional CI), then delayed histone deposition, improper chromosome condensation and cell division abnormalities result in embryonic mortality (Landmann, et al. 2009; Lassy and Karr 1996; Serbus, et al. 2008; Tram and Sullivan 2002). Other described reproductive manipulations include parthenogenesis (Stouthamer, et al. 1990), male-killing (Hurst, et al. 1999), and feminization (Rousset, et al. 1992), all of which give a fitness advantage to *Wolbachia*-infected females and thus assist the spread of the infected matriline through a population. These manipulations, once sustained, can also impact host evolution including speciation (Bordenstein, et al. 2001; Brucker and Bordenstein 2013; Jaenike, et al. 2006) and mating behaviors (Miller, et al. 2010; Moreau, et al. 2001; Randerson, et al. 2000; Shropshire and Bordenstein 2016).

In addition to the aforementioned reproductive manipulations, *Wolbachia* strains affect host biology by provisioning nutrients (Hosokawa, et al. 2010), altering host survivorship (Min and Benzer 1997) and fecundity (Dedeine, et al. 2001; Stouthamer and Luck 1993), and importantly, protecting the host against pathogens (Bian, et al. 2010; Hughes, et al. 2011; Kambris, et al. 2009; Moreira, et al. 2009; Teixeira, et al. 2008; Walker, et al. 2011). The combination of reproductive manipulations that enable *Wolbachia* to spread in a population, and the ability to reduce vector competence through pathogen protection, have placed *Wolbachia* in the forefront of efforts to control target arthropod populations (Bourtzis, et al. 2014; Hoffmann, et al. 2011; Turelli and Hoffmann 1991; Walker, et al. 2011; Zabalou, et al. 2004). Despite these important applications, the widespread prevalence of *Wolbachia* across arthropod taxa (Hilgenboecker, et al. 2008; Werren and Windsor 2000; Zug and Hammerstein 2012), and decades of research, only recently have the genes underlying CI been determined (Beckmann, et al. 2017; LePage, et al. 2017).

Two studies converged on the same central finding: coexpression of a pair of syntenic genes recapitulates the CI phenotype (Beckmann, et al. 2017; LePage, et al. 2017). Uninfected *Drosophila melanogaster* males transgencially expressing the two genes from *w*Mel *Wolbachia* caused CI-like embryonic lethality when crossed with uninfected females that was notably rescued by *w*Mel-infected females (LePage, et al. 2017). Additionally, the two *w*Mel genes separately enhanced *Wolbachia*-induced CI in a dose dependent manner when expressed in *Wolbachia*-infected males (LePage, et al. 2017). In a separate study, CI-like embryonic lethality was also recapitulated through transgenic coexpression in *D. melanogaster* males of homologous transgenes encoded by the *Wolbachia w*Pip strain (which infects *Culex* mosquitoes) (Beckmann, et al. 2017). These two genes occur in the recently discovered eukaryotic association module of temperate phage WO (Bordenstein and Bordenstein 2016), which was previously implicated in influencing CI (Bordenstein, et al. 2006; Duron, et al. 2006; Masui, et al. 2000; Sinkins, et al. 2005). The presence of these genes within prophage WO has implications for the transmission of these genes, namely vertical transmission in the *Wolbachia* genome versus horizontal transfer of phage WO. The genes were proposed as candidate CI effectors due to the presence of one of the protein products in the spermathecae of infected female mosquitoes (Beckmann and Fallon 2013) and their absence in the *w*Au *Wolbachia* strain that lost CI function (Sutton, et al. 2014).

The *w*Mel homologs of these genes are designated cytoplasmic incompatibility factors *cifA* (locus WD0631) and *cifB* (locus WD0632), with *cifA* always encoded directly upstream of *cifB* (LePage, et al. 2017). The gene set occurrs in varying copy number across eleven total CI-inducing strains that correlates with CI levels. Core sequence changes of the two genes exhibit a pattern of codivergence and in turn closely match bidirectional incompatibility patterns between *Wolbachia* strains. Homologs of CifA and CifB protein sequences belong to four distinct phylogenetic types (designated Types I – IV) that do not correlate with various phylogenies of *Wolbachia* housekeeping genes or phage WO *gpW* (locus WD0640) (LePage, et al. 2017). The homologous sequences in *w*Pip also cluster in Type I, though they are 66% and 76% different from *w*Mel’s, respectively (Beckmann, et al. 2017). Hereinafter we use *cifA* and *cifB* to refer to these genes, unless specifically referring to analyses of the *w*Pip homologs, *cidA* and *cidB*. *In vitro* functional analyses revealed that *cidB* can encode deubiquitylase activity, and *cidA* encodes a protein that binds CidB (Beckmann, et al. 2017). Mutating the catalytic residue in the deubiquitylating domain of CidB results in a loss of the CI-like function in transgenic flies (Beckmann, et al. 2017). Whether these genes have additional enzymatic or regulatory roles and which other residues are important for function remain open questions.

There are important considerations for the location, organization, and characterization of these genes. Whether or not *cifA* and *cifB* form a strict, toxin-antitoxin operon is debatable, and likewise has important implications for how gene expression is regulated by *Wolbachia* during host infection. Support for the operon hypothesis is based on weak transcription across the junction between *cidA* and *cidB*, inferred to be due to the presence of polycistronic mRNA (Beckmann and Fallon 2013; Beckmann, et al. 2017); an alternative explanation is transcriptional slippage. Quantitative transcription analyses and computational predictions of operon structure do not support the operon hypothesis (LePage, et al. 2017). Moreover and importantly, transgenic studies show that both *cifA* and *cifB* are required for induction of CI and thus cannot form a strict toxin (*cifB*) - antitoxin (*cifA*) system. As both genes encode CI function and can individually enhance *Wolbachia*-induced CI, and there is mixed evidence for classification as an operon, it does not appear that characterization as a strict toxin-antitoxin operon is warranted (LePage, et al. 2017). However, like toxin-antitoxin systems, CidA binds CidB *in vitro* and expression of *cidA* rescues temperature-sensitive growth inhibition induced by *cidB* expression in *Saccharomyces*, via an as-yet-unknown mechanism (Beckmann, et al. 2017).

As it stands now, the genes remain largely unannotated with the exception of a few small domains. If other predicted protein domains occur in *cifA* and *cifB*, they would provide new hypotheses for the mechanism of CI. Finally, the sequence diversity and/or loss of *cif* genes across the *Wolbachia* tree may give insights into the selective conditions that maintain the *cif* genes versus those that do not. Exploration of *cif* gene regulation, expression, and function thus can provide a framework for more targeted investigations of *Wolbachia*-host interactions, and potentially inform the deployment of *Wolbachia*-based arthropod control.

## Materials and Methods

### Expression

For analysis of RNAseq data we used our published approach (Gutzwiller, et al. 2015). Briefly, fastq sequences for 1 day old male and female flies were mapped against the *Wolbachia w*Mel reference genome (Ensembl Genomes Release 24, Wolbachia_endosymbiont_of_drosophila_melanogaster.GCA_000008025.1.24) using bwa mem v. 0.7.5a with default parameters in paired-end mode. Mapped reads were sorted and converted to BAM format using samtools v0.1.19 after which BAM files were used as input to Bedtools (bedcov) to generate pileups and count coverage at each position. For expression correlations between genes, the raw RNAseq counts were divided by (gene length + 99), where 99 corresponds to read length (100) – 1. Within a growth stage these values were multiplied by 1e^6^ / (sum of values in stage) (Li and Dewey 2011). A pairwise distance between all genes was defined as (1 – R), where the R is the Pearson correlation coefficient between the normalized expression values of two genes. Possible negative correlations would be “penalized” here, resulting in a larger distance. Distances were clustered using the Kitsch program of PHYLIP (Felsenstein 1989).

### Operon Prediction *in silico*

We used the dynamic profile of the transcriptome above to identify operons within the *w*Mel genome using two different approaches. We used the program Rockhopper (McClure, et al. 2013), using default parameters, in conjunction with the BAM files generated above to delineate likely operons across the entire genome. In addition, we took a fine-scale approach, focusing on the junction between *cifA* and *cifB* (Fortino, et al. 2014), using the pileup files generated above and identifying drops in gene expression correlated to genomic position using a sliding window analysis.

### Nucleic Acid Extractions and Quantitative PCR

To identify *Wolbachia* gene expression in adult male and female *D. melanogaster*, RNA was extracted from individual, age-matched flies (1-3 days old, stock 145) using a modified Trizol extraction protocol. Briefly, 500 uL of Trizol was added to individual flies and samples homogenized using a pestle. After a 5-minute incubation at room temperature, a 12,000 rcf centrifugation (at 4C for 10 min) was followed by a chloroform extraction. Aqueous phase containing RNA was extracted a second time with phenol:chloroform before isopropanol precipitation of RNA. This RNA pellet was washed and resuspended in THE RNA Storage Solution (Ambion). To detect the number of *cifA* and *cifB* transcripts as well as RNA levels across the junction between *cifA* and *cifB*, we utilized the RNA extracted from these flies and the SensiFAST SYBER Hi-ROX One-step RT mix (Bioline) and the Applied Biosystems StepOne Real-time PCR system with the following primer sets: *cifAF*: ATAAAGGCGTTTCAGCAGGA, *cifAR*: TCAATGAGGCGCTTCTAGGT; *cifBF*: TACGGGAAGTTTCATGCACA, *cifBR*: TTGCCAGCCATCATTCATAA; *cifA*_endF: TCTGGTTCTCATAAGAAAAGAAGAATC, *cifB*_begR: AACCATCAAGATCTCCATCCA. As a reference for transcription activity of the core *Wolbachia* genome, we utilized the *Wolbachia ftsZ* gene (Forward: TTTTGTTGTCGCAAATACCG; Reverse: AGCAAAGCGTTCACATTTCC). We designed primers to *ftsZ* because as a core protein involved in cell division, the quantities of *ftsZ* would better correlate with bacterial numbers and activity. Reactions were performed in duplicate or triplicate in a 96-well plate and CT values generated by the machine, were used to calculate the relative amounts of *Wolbachia* using the ΔΔCt (Livak) method.

### Correlated Cif Trees and Distance Matrices

Quantifying congruence scores between the CifA and CifB trees was carried out with Matching Cluster (MC) and Robinson Foulds (RF) metrics using a custom python script previously described (Brooks, et al. 2016) and the TreeCmp program (Bogdanowicz, et al. 2012). MC weights topological congruency of trees, similar to the widely used RF metric. However, MC takes into account sections of subtree congruence and therefore is a more refined evaluation of small topological changes that affect incongruence. Significance in the MC and RF analyses was determined by the probability of 100,000 randomized bifurcating dendrogram topologies yielding equivalent or more congruent trees than the actual tree. Normalized scores were calculated as the MC and RF congruency score of the two topologies divided by the maximum congruency score obtained from random topologies. The number of trees that had an equivalent or better score than the actual tree was used to calculate the significance of observing that topology. Mantel tests were also performed on the CifA and CifB patristic distance matrices calculated in Geneious v8.1.9 (Kearse, et al. 2012). A custom Jupyter notebook (Pérez and Granger 2007) running python v3.5.2 (http://python.org) was written in the QIIME2 (Caporaso, et al. 2010) anaconda environment, and the Mantel test (Mantel 1967) utilized the scikit-bio v0.5.1 (scikit-bio.org) Mantel function run using scikit-bio distance matrix objects for each gene. The Mantel test was run with 100,000 permutations to calculate significance of the Pearson correlation coefficient between the two matrices using a two-sided correlation hypothesis.

### Genomes Used in Comparative Analyses

In order to identify *cif* homologs across the *Wolbachia* genomes, we defined orthologs across existing, sequenced genomes using reciprocal best blastp. We included *Wolbachia* genomes across five Supergroups: monophyletic clades of *Wolbachia* based on housekeeping genes, denoted by uppercase letters (O’Neill, et al. 1992; Werren, et al. 1995). Supergroups A and B are the major arthropod infecting lineages, while C and D infect nematodes (Bandi, et al. 1998). Supergroup F *Wolbachia* infect a variety of hosts (Lo, et al. 2002). Included in this analysis were 11 type A strains (*w*Ri, *w*Ana, wSuzi, *w*Ha, *w*Mel, *w*MelPop, *w*Au, *w*Rec, *w*Gmm, *w*Uni, *w*VitA), 10 type B strains (*w*PipJHB, *w*PipPel, *w*PipMol, *w*Bol1-b, *w*Bru, *w*CauB, *w*No, *w*Tpre, *w*AlbB, *w*Di), 2 type C strains (*w*Ov, *w*Oo), and one each type D (*w*Bm) and type F (*w*Cle). We included all genomic data available for each strain such that if multiple assemblies existed for each *Wolbachia* variant (such as in the case of *w*Uni) we included the union of all available contigs for that strain. *Wolbachia* orthologs were defined based on reciprocal best blast hits between amino acid sequences in *Wolbachia* genomes. An orthologous group of genes was defined by complete linkage such that all members of the group had to be the reciprocal best hit of all other members of the group. *w*Ana, *w*Gmm, *w*PipMol, *w*Bru, and *w*CauB were not used in subsequent analyses due to problematic assemblies. Information on strain phenotypes, hosts, and accession numbers can be found in Table 1.

**Table 1.**
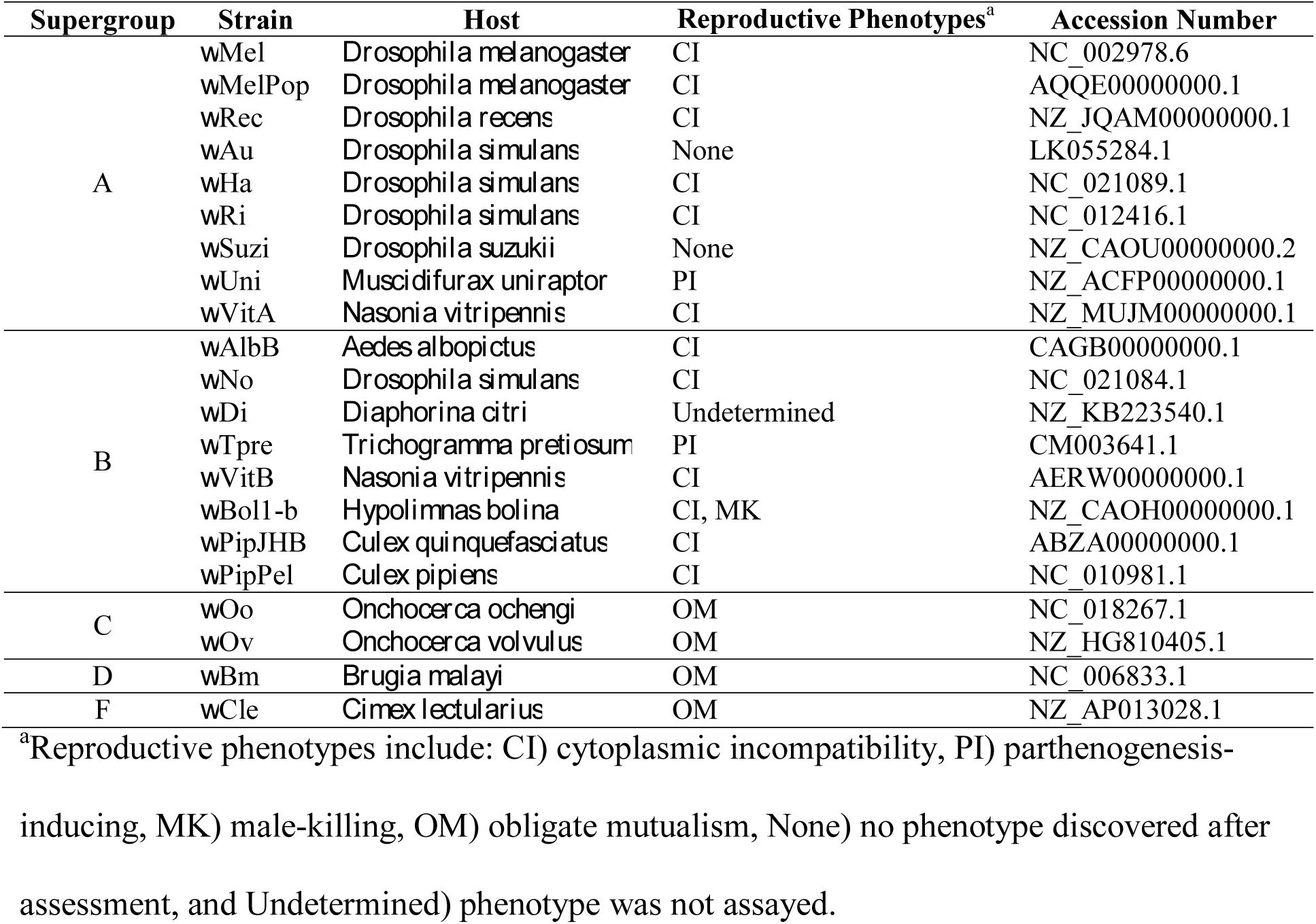
Strains used in comparative analyses of *cifA* and *cifB*.

### Cif Phylogenetics

CifA and CifB protein sequences were identified using BLASTp searches of WOMelB WD0631 (NCBI accession number AAS14330.1) and WD0632 (AAS14331.1), respectively. Homologs were selected based on: 1) *E*☐=☐≤☐10^−30^, 2) query coverage greater than 70%, and 3) presence in fully sequenced *Wolbachia* genomes. All sequences were intact with the exception of a partial WOSuziC CifA (WP_044471252.1) protein. The missing N-terminus was translated from the end of contig accession number CAOU02000024.1 and concatenated with partial protein WP_044471252.1 for analyses, resulting in 100% amino acid identity to WORiC CifA (WP_012673228.1). In addition, two previously identified sequences (LePage, et al. 2017), WORecB CifB and WORiB CifB, were not available in NCBI’s database and translated from nucleotide accession numbers JQAM01000018.1 and CP001391.1, respectively. The previously identified WOSol homologs (CifA: AGK87106 and CifB: AGK87078) (LePage, et al. 2017) were also included in our analyses. All protein sequences were aligned with the MUSCLE (Edgar 2004) plugin in Geneious Pro version 8.1.7 (Kearse, et al. 2012); the best models of evolution, according to corrected Akaike (Hurvich and Tsai 1993) information criteria, were estimated to be JTT-G using the ProtTest server (Abascal, et al. 2005); and phylogenetic trees were built using the MrBayes (Ronquist, et al. 2012) plugin in Geneious.

### Protein Structure

All candidate CI gene protein sequences were assessed for the presence of domain structure using HHpred (https://toolkit.tuebingen.mpg.de/hhpred/ (Söding, et al. 2005)) with default parameters and the following databases: SCOPe95_2.06, SCOPe70_2.06, cdd_04Jul16, pfamA_30.0, smart_04Jul16, COG_04Jul16, KOG_04_Jul16, pfam_04Jul16, and cd_04Jul16. Schematics were created in inkscape (https://inkscape.org/), to show regions with significant structural hits, at a corrected p-value of p < 0.05. Modules were defined based on the presence of multiple highly significant hits within a region.

### Protein Conservation

Protein conservation was determined with the Protein Residue Conservation Prediction tool (http://compbio.cs.princeton.edu/conservation/index.html (Capra and Singh 2007)), using aligned amino acid sequences, Shannon entropy scores, a window size of zero, and sequence weighting set to “false”. Conservation was subsequently plotted in R version 3.3.2, and module regions were delineated according to coordinates of the WOMelB modules within the alignment. CI gene conservation scores were calculated separately for Type I sequences, and for all types together. For CifB Type I sequences, the WOVitA4 ortholog was left out, due to the extended C-terminus of that protein. Conservation scores were also calculated for “control proteins”: Wsp (*Wolbachia* surface protein), known to be affected by frequent recombination events (Baldo, et al. 2005), and FtsZ, which is relatively unaffected by recombination (Baldo, et al. 2006b; Ros, et al. 2009). Variation in amino acid conservation between modules and non-module regions was assessed in R version 3.3.2 with a one-way ANOVA including “region” (either the unique module number, or “non-module”) as a fixed effect, and followed by Tukey Honest Significant Difference for post hoc testing.

### Cif Modules

The WOMelB structural regions delineated by HHpred were used to search for the presence of Cifs or remnants of Cifs across the *Wolbachia* phylogeny. Amino acid sequences of the WOMelB modules were queried against complete genome sequences (Table 1) using tblastn. Any hit that was at least 50% of the length and 30% identity, or at least 90% of the length and 20% identity of the WOMelB module was considered a positive match. Module presence was plotted across a *Wolbachia* phylogeny constructed using the five Multi Locus Sequence Typing (MLST) genes defined by Baldo *et al*. (Baldo, et al. 2006b). Nucleotide sequences were aligned with MAFFT version 7.271 (Katoh and Standley 2013), and concatenated prior to phylogenetic reconstruction with RAxML version 8.2.8 (Stamatakis 2014), the GTRGAMMA substitution model, and 1000 bootstrap replicates.

### Hidden Markov Model Searches

To identify *cif* homologs in draft *Wolbachia* genome assemblies we used the program suite HMMER (Eddy 2011). We defined *cif* types based on our phylogenetic trees (Figure 4) and used aligned amino acids from these types as input to HMMBUILD, using default parameters. We then searched six *Wolbachia* WGS assemblies (NCBI project numbers PRJNA310358, PRJNA279175, PRJNA322628) using HMMSEARCH with –F3 1e-20 –cut_nc and –domE 1e-10. Regardless of thresholds used, or *cif* type of HMM, resulting hits did not differ.

## Results

### *cifA* and *cifB* are Not Co-transcribed or Co-regulated and Do Not Comprise an Operon in *w*Mel

To assess the operon hypothesis, we reasoned that genes which are co-transcribed and co-regulated will exhibit the following properties: similar total expression levels in whole animals and correlated gene expression across host development. We therefore utilized an existing RNAseq dataset for *Wolbachia* in *Drosophila melanogaster*, covering 24 life cycle stages and 3 time samplings each for adult males and females (Gutzwiller, et al. 2015). We mapped reads to the existing *w*Mel assembly (see methods), and calculated Pearson correlation coefficients for normalized expression values for each pairwise comparison across host development. In adult males and females, *cifA* and *cifB* in *w*Mel are not expressed at similar levels (Figure 1), with *cifA* expressed at significantly higher levels compared to *cifB* (eight-fold higher based on RPKM values across both genes).

**Fig 1.**
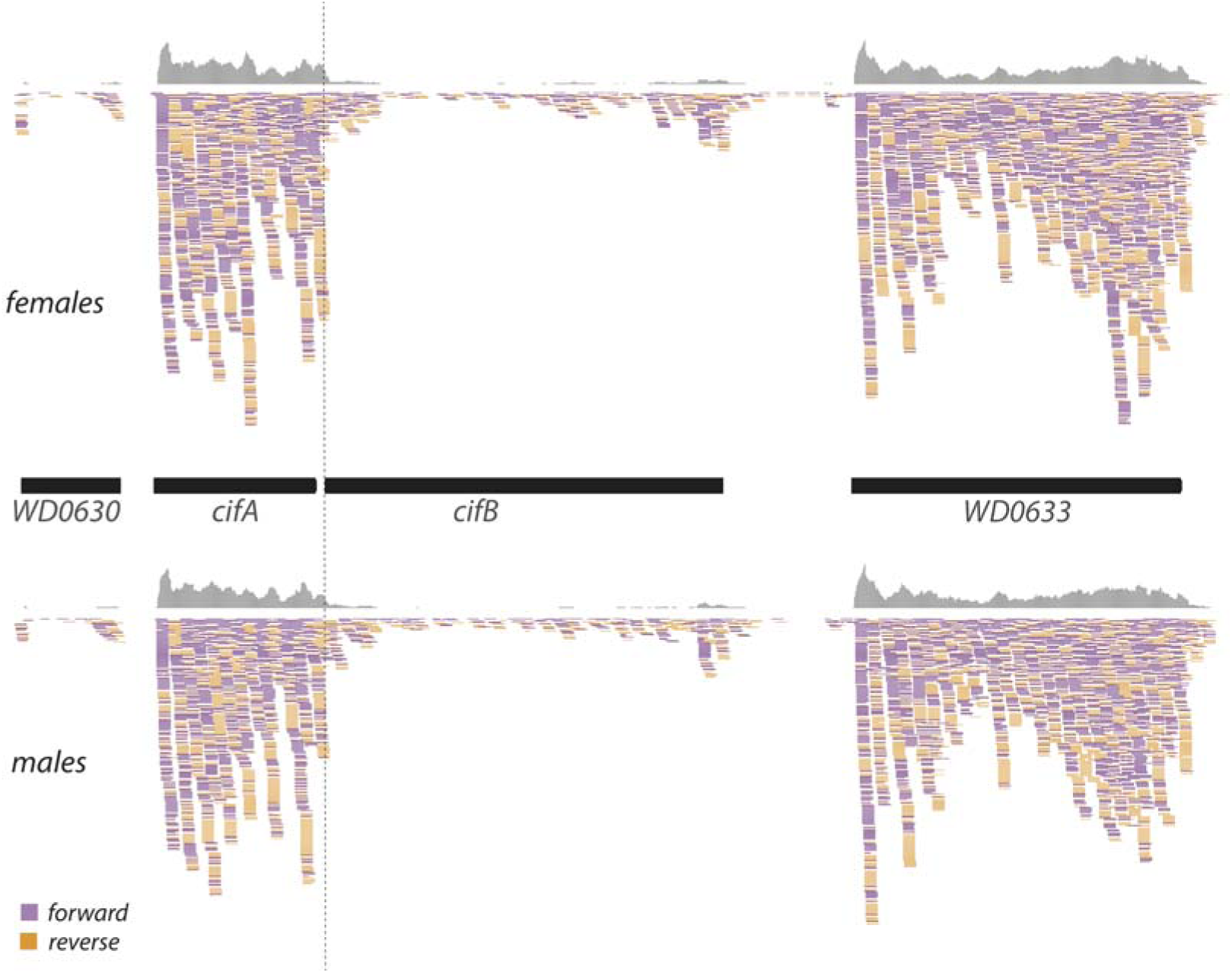
RNASeq analysis of *cifA* and *cifB* gene expression in whole adult, 1 day old female and male *Drosophila melanogaster* flies. Raw reads were mapped to the *w*Mel assembly (using bwa) and coverage visualized using the Integrated Genomics Viewer (v2.3.77). The start of the *cifB* open reading frame is denoted by a vertical, dotted line.

To further explore expression of the *cif* genes in *w*Mel and assess whether or not polycistronic mRNA is produced, we performed a quantitative PCR analysis of gene expression from three-day old male and female flies (Figure 2). We observed transcripts covering the junction between *cifA* and *cifB*. However, transcripts covering this junction were much more similar to expression levels in *cifA*, while expression of *cifB* was nine-fold less. Therefore, as *cifA* and *cifB* are separated by only 76 bp, distinguishing between 3’ UTRs from *cifA* and full *cifA*-*cifB* transcripts is not possible.

**Fig 2.**
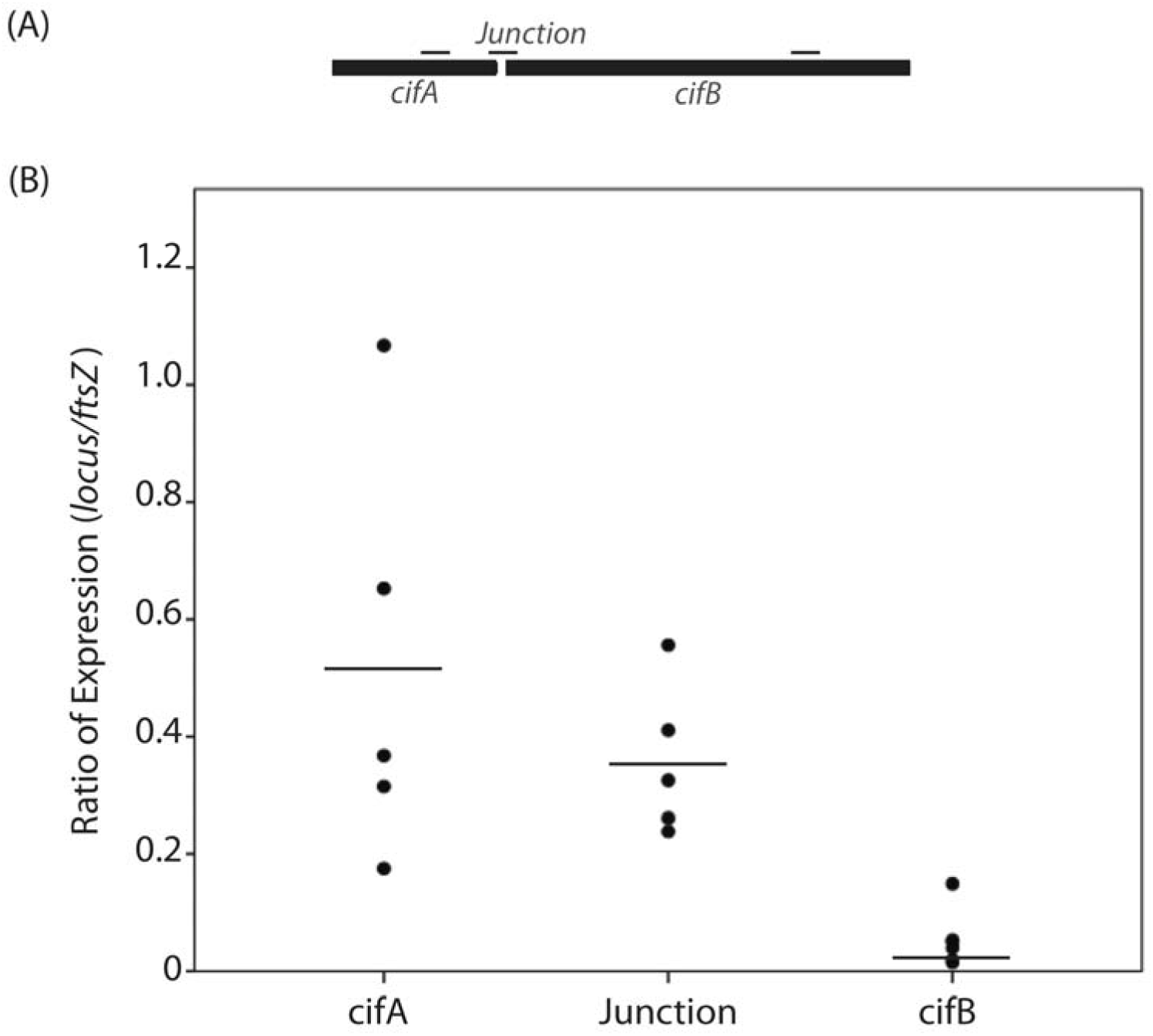
Relative expression ratio of *cifA*, the junction between *cifA/cifB*, and *cifB* to *ftsZ*. Expression of both genes and their junction was quantified using qRT-PCR, and normalized to *Wolbachia ftsZ* gene expression. *cifB* gene expression is significantly less than that of the junction (t= 3.220, df=16, p=0.005) and less than *cifA* (t=-3.840, df=17, p=0.001).

We next used two computational methods to test for a potential operon between *cifA* and *cifB* using our RNAseq analyses. After mapping reads to the *w*Mel assembly, we used the resulting BAM files as input to Rockhopper (McClure, et al. 2013). The program was able to correctly identify known operons in *w*Mel (such as the T4SS WD0004-WD0008 and the ribosomal protein operon) but it did not identify *cifA* and *cifB* as an operon. We also used a sliding-window approach, using pileup files generated as part of the mapping, to identify correlations between genomic position and gene expression drops in the RNAseq data, as in (Fortino, et al. 2014).The two open reading frames for *cifA* and *cifB* span positions 617223-618647 and 618723-622223, respectively. From positions 618600 to 618700, we observe a significant positive correlation between coverage and genomic location (Pearson Correlation = 0.99, p < 0.001). However, across the junction between *cifA* and *cifB* (position 618700), we saw a very large drop in gene expression in both males and females (from an average coverage of 4616 to 38 per position). This result suggests that *cifA* and *cifB* are not co-transcribed.

Finally, we clustered the *w*Mel *cif* genes based on their similarity in expression across *Drosophila* development (Supplemental Figure S1). *cifA* did not group with *cifB* in *w*Mel (Figure 3), suggesting that these two genes are not co-regulated. Indeed, the pattern of *cifA* expression differs strikingly from that of *cifB*. *cifB* is expressed during embryogenesis and generally down-regulated in pupae and adults, while *cifA* is highly expressed in adult males and females and late time points during embryogenesis (Figure 1). Curiously, the expression profile of *cifA* in flies during development is most closely correlated with the *wsp* locus WD1063 (Figure 3).

**Fig 3.**
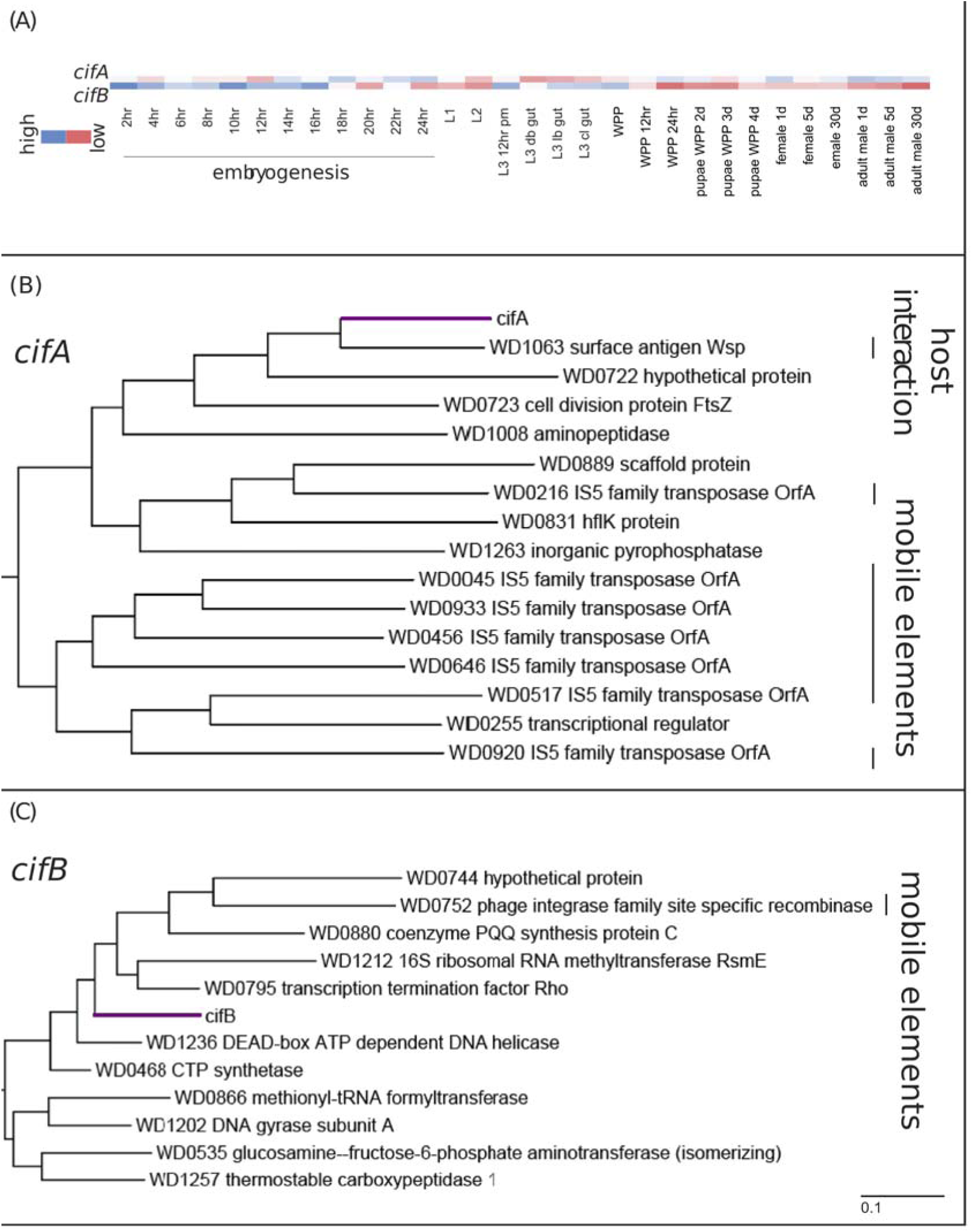
Gene expression of *cifA* and *cifB* during *Drosophila melanogaster* development. A) Heatmap representation of normalized transcripts per kilobase million (TPM) for both *cifA* and *cifB* during *Drosophila melanogaster* development. *cifB* is highly expressed during embryogenesis and downregulated after pupation while *cifA* is more highly expressed in adults and pupae. Clustering of *Wolbachia* loci based on expression across fly development illustrates correlated expression profiles between *w*Mel loci and *cifA* (B) or *cifB* (C). Mobile elements and loci involved in host interaction (*wsp*) are indicated with vertical lines on the right side of the figure.

### New Protein Domain Predictions are Variable Across the Cif Phylogeny

We recovered the four previously identified phylogenetic types (LePage, et al. 2017). Here, our analyses include additional strains that cause reproductive parasitism beyond CI (parthenogenesis and male-killing, Table 1), and the more divergent Type IV paralogs for *cifA*, so far identified in B-Supergroup *Wolbachia*. We recover a set of Type III alleles from *w*Uni, a strain that induces parthenogenesis in the parasitoid wasp, *Muscidifurax uniraptor* (Stouthamer, et al. 1993). The *w*Bol1-b strain, a male-killer that has retained CI capabilities (Hornett, et al. 2008), has alleles belonging to both Type I and Type IV.

Homologs and predicted protein domains of CifA and CifB for all four phylogenetic types (LePage, et al. 2017) from *Wolbachia* strains that cause CI, parthenogenesis, male-killing, or no reproductive phenotype were characterized by HHpred homology and domain structure prediction software (Söding, et al. 2005). Search parameters are described in the methods. Several new prominent protein domains (as determined by the presence of multiple highly significant structural predictions within a region), herein referred to as “modules”, were identified for each CifA and CifB protein sequence. In Table 2 we list the prominent module annotations identified across CifA and CifB Types. Multiple structural hits within a region can be explained by the homology of the significant domains predictions to each other.

**Table 2.**
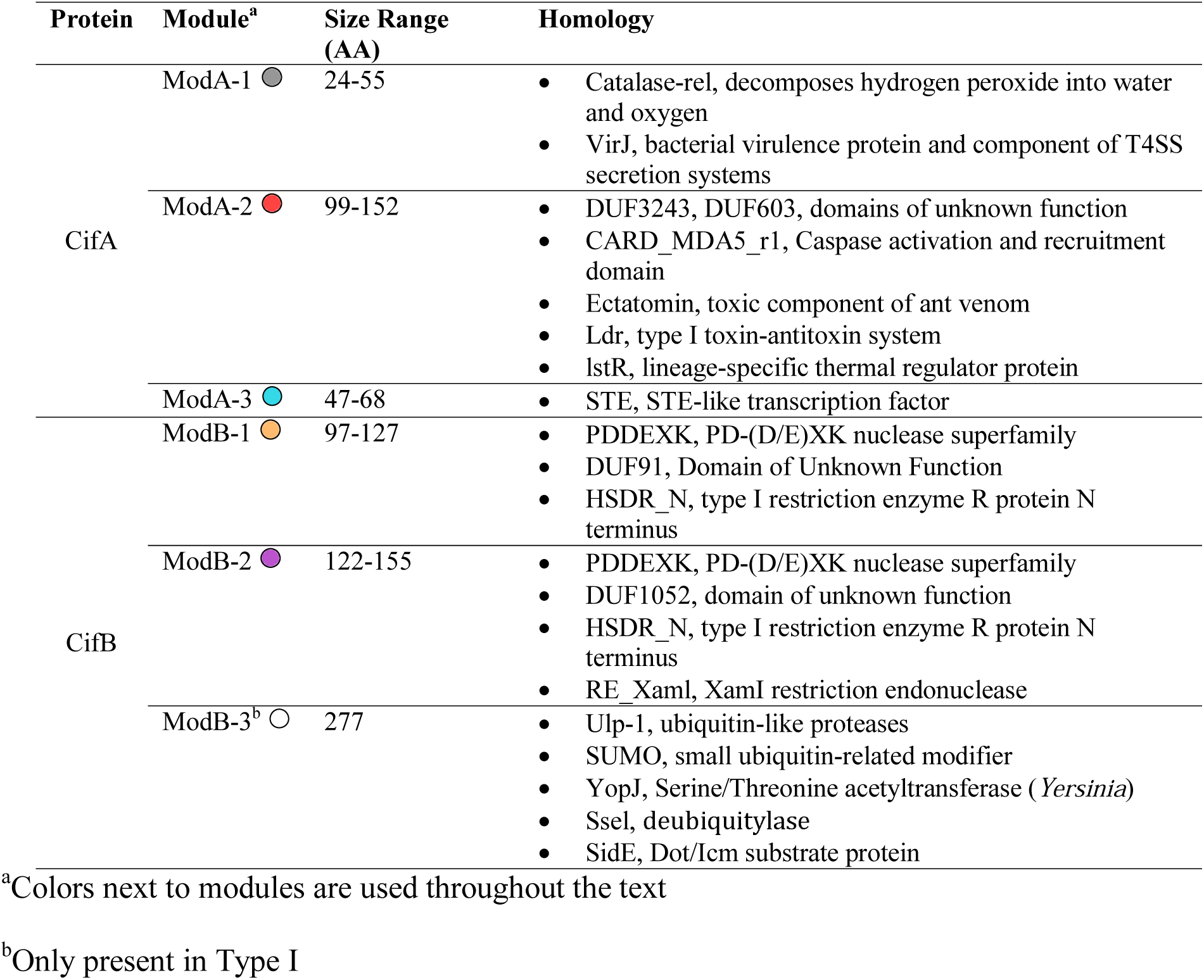
Predicted structural modules of Cif proteins.

For CifA, three main modules were annotated (Figure 4A, Table 2). First, the most N-terminal module (ModA-1) in Type I, II and III variants shows homology to Catalase-rel (p = 0.001-0.003), which is predicted to catalyze the breakdown of hydrogen peroxide (Chelikani, et al. 2004) (Type I) and protect the cell from toxic effects, or VirJ (p = 0.002-0.003), a bacterial virulence protein and component of T4SS secretion systems (Pantoja, et al. 2002) (Types II and III). The second CifA module in the central region (ModA-2) has homology to a caspase recruitment domain (p = 0.005-0.009), venom and toxin-related domains (p ≤ 0.001), and a thermal regulator protein (p = 0.002). The very significant homology to a toxin is interesting, given that CifA was hypothesized to act as an antitoxin. Notably, CifA is required for and enhances the induction of CI (LePage, et al. 2017), which contradicts its proposed function as simply an antitoxin (Beckmann, et al. 2017). The last CifA module in the C-terminal region (ModA-3) has multiple strong hits to a STE-like transcription factor (p ≤ 0.001). There were additional annotations that emerged due to weak or singular matches. In Type IV variants, there is a separate N-terminal region that shares homology with a conserved eukaryotic family with potential methyltransferase activity, FAM86 (p = 0.003). Most Type I alleles have C-terminal homology to a nuclear cap-binding protein that binds RNA (p = 0.010 – 0.020). WOHa1, WOBol1b, and WOSol have an additional N-terminal region containing a conserved domain of unknown function (p = <0.001 – 0.005). Type IV genes have a yeast-like salt tolerance down-regulator domain NST1 (p = 0.003). Lastly, WOVit4 and *w*Uni lack the most N-terminal CifA homology region, ModA-1.

**Fig 4.**
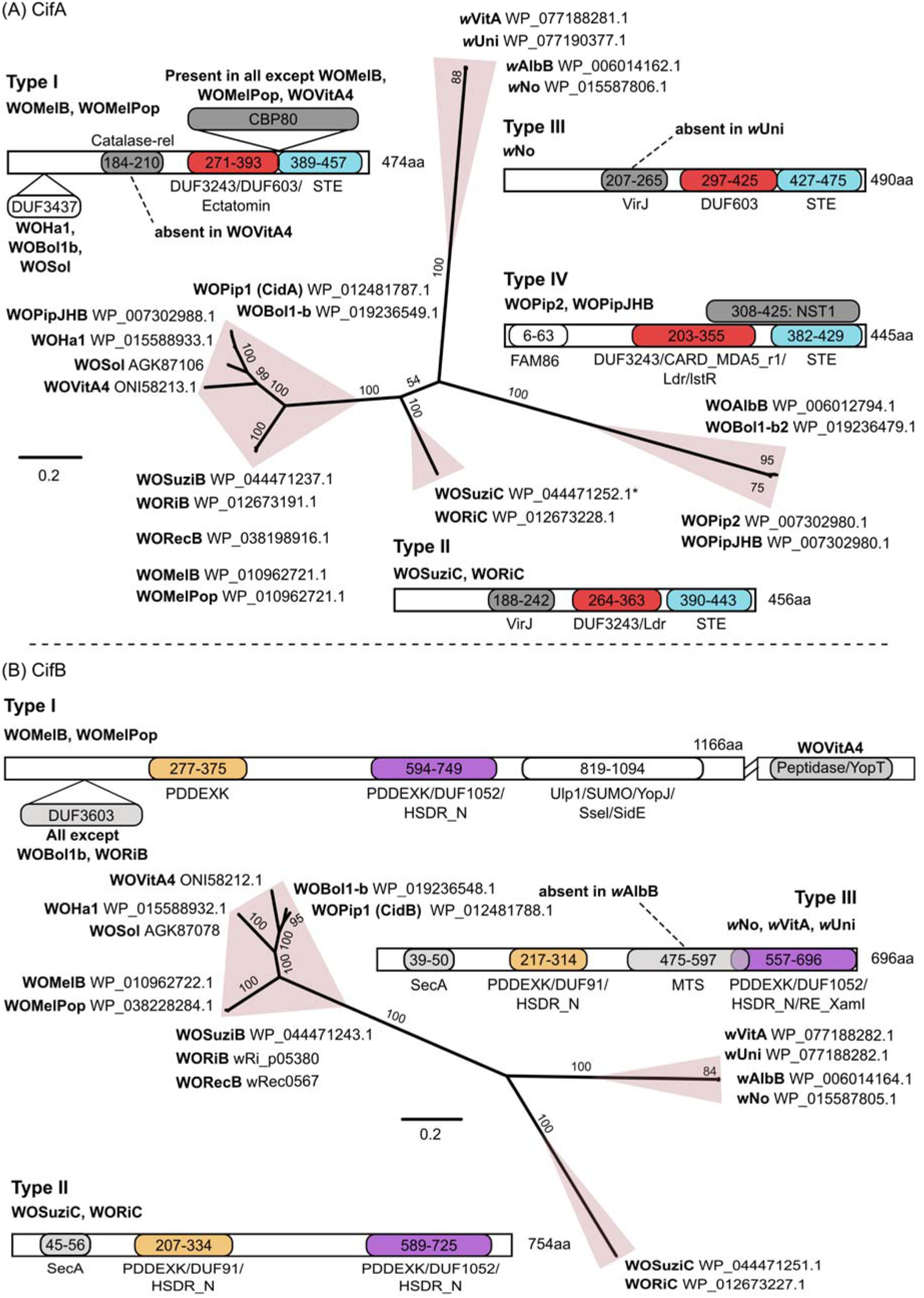
Phylogenetic relationships and representative predicted protein structure of Cif protein types. A) CifA and B) CifB. Alleles are in bold next to their corresponding accession number, and pink shapes around branches designate monophyletic “types”. Representative structures are shown for each type, with the length of the protein indicated at the C-terminus. Variations in within-type structure are shown. If an allele is not listed as a representative, and significant structural variations are not indicated, then only the exact coordinates of the structural regions differed by a few amino acids. All HHpred structural predictions are significant at a corrected p-value of < 0.05, and listed in order of ascending p-value for regions with multiple structural hits. Allele names use the previously described naming convention with a WO prefix referring to particular phage haplotype, and the *w* prefix indicating a phage-like island (LePage, et al. 2017). The N-terminus of WOSuziC (*) was translated from the end of another contig and concatenated to get the full-length protein (see methods). WOMelB and WOMelPop are identical at the amino acid level, as are WOPipJHB and WOPip2.

For CifB, three main modules were defined (Figure 4B, Table 2). The first (ModB-1) and second (ModB-2) most N-terminal regions both have matches to the PDDEXK nuclease family (p < 0.001), the HSDR_N restriction enzyme (p = < 0.001-0.010), and domains of unknown function (DUF1052, DUF91). The third module, found only in the Type I C-terminus (ModB-3), has very strong homology to a number of ubiquitin-modification and peptidase domains (p < 0.001), as well as YopJ, which in *Yersinia,* aids in infecting a eukaryotic host (Paquette, et al. 2012) (p = < 0.001-0.020). ModB-3 contains the catalytic residue associated with toxicity/CI function in CidB (Beckmann, et al. 2017). In addition to the annotated modules, all Type I alleles except WOBol1b and WORiB have a single hit to a conserved domain of unknown function in the N-terminus (p = 0.001 – 0.005), and Type III alleles (except for *w*AlbB) have a region of homology to a methyltransferase domain (MTS) (p < 0.001). Both Type II and III alleles have a single short hit in the N-terminus to a SecA regulator. WOVitA4 (Type 1) has an extended C-terminus not present in any other alleles, and within that extended C-terminus is an additional peptidase/YopT-like region, similar to ModB-3. CifB Type IV alleles (WOAlbB, WOPip2, and *w*Bol1-b) were not included in the phylogenetic reconstruction, as they are highly divergent and not reciprocal blasts of WOMelB *cifB*. Despite their divergence, these Type IV CifB alleles have similar structures to Type II and III alleles: two PDDEXK-like modules, and no Ulp-1-like module three (Supplemental Figure S3). Full structural schematics with exact coordinates and homology regions for each allele are available in the supplemental material (Supplemental Figures S2 and S3), as are all significant domain hits with associated p-values and extended descriptions (Supplemental Tables S1 and S2).

### CifA and CifB Codiverge

Initial phylogenetic trees based on core amino acid sequences of Type I-III variants of CifA and CifB exhibited similar trees (LePage, et al. 2017). Here we statistically ground the inference of codivergence using the largest set of *Wolbachia* homologs to date. We quantified congruence between the CifA and CifB phylogenetic trees for Types I-III (Supplemental File S1) using Matching Cluster (MC) and Robinson–Foulds (RF) tree metrics (Bogdanowicz and Giaro 2013; Bogdanowicz, et al. 2012; Robinson and Foulds 1981), with normalized distances ranging from 0.0 (complete congruence) to 1.0 (complete incongruence). Results show strong levels of congruence between CifA and CifB (p < 0.00001 for both, normalized MC = 0.06 and normalized RF = 0.125). To further statistically validate the inference of codivergence, we measured the correlation between patristic distance matrices for CifA and CifB using the Mantel test (Mantel 1967). Results demonstrate a high degree of correlation between patristic distance matrices, and through permutation show that independent evolution of CifA and CifB is highly unlikely (Pearson correlation coefficient = 0.905, p = 0.00001).

### Cif Proteins Evolve Rapidly

Amino acid sequence conservation across the full length of the Cif proteins was determined and compared to *Wolbachia* amino acid sequences of genes that either have signatures of recombination and directional section (Wsp, *Wolbachia* surface protein) or have not undergone extensive recombination and directional selection (FtsZ, cell division protein). Wsp protein sequences exhibit considerable divergence (mean conservation = 0.85), with very few sites in a row being completely conserved (Figure 5A). In contrast, FtsZ is relatively conserved (mean conservation = 0.94), and most of the divergence is clustered at the C-terminus (Figure 5B). Mean conservation for the Cif protein sequences were lower than Wsp - 0.83 for Type I CifA alleles (Figure 5C) and 0.82 for Type I CifB alleles (Figure 5E, Table 3). When all Cif alleles were considered, mean conservation was even further reduced - 0.58 for CifA (Figure 5D) and 0.43 for CifB (Figure 5F). The lower average conservation of CifB genes is in part due to the many insertions and deletions in the alignment, and the missing C-terminal deubiquitylase region, ModB-3, of the Type II and III alleles. Thus, several CifB proteins apparently lack this activity, and whether these variants cause CI remains to be determined. Importantly, although the CifB proteins are highly divergent, the catalytic residue (red dot in Figures 5E and 5F) in the deubiquitylating module of CifB is unique to and completely conserved for the Type I alleles. The Cif proteins have extensive amounts of diversity, with completely conserved amino acids distributed across the length of the protein, and not confined to any particular regions (Figure 5C-F, Supplemental Tables S3-S6). There were significant differences in the level of conservation between modules and non-module regions for the Type I alignments of both CifA (F_3,495_ = 11.75, p = 0.0021) and CifB (F_3,1195_ = 11.75, p = 1.38e-07) (Table 3). The only module that had significantly higher conservation than the non-module regions of the alignment was ModB-1 (p = 0.0173). The *w*Mel strain contains the (P)D-(D/E)XK motif (blue dots in Figures 5E and 5F) (Kosinski, et al. 2005), but it is less than 80% conserved across strains despite the higher average conservation of this module. In contrast, ModA-3 and ModB-3 are significantly less conserved than the non-module regions of the corresponding proteins (CifA, p = 0.0400; CifB, p = 0.0001).

**Fig 5.**
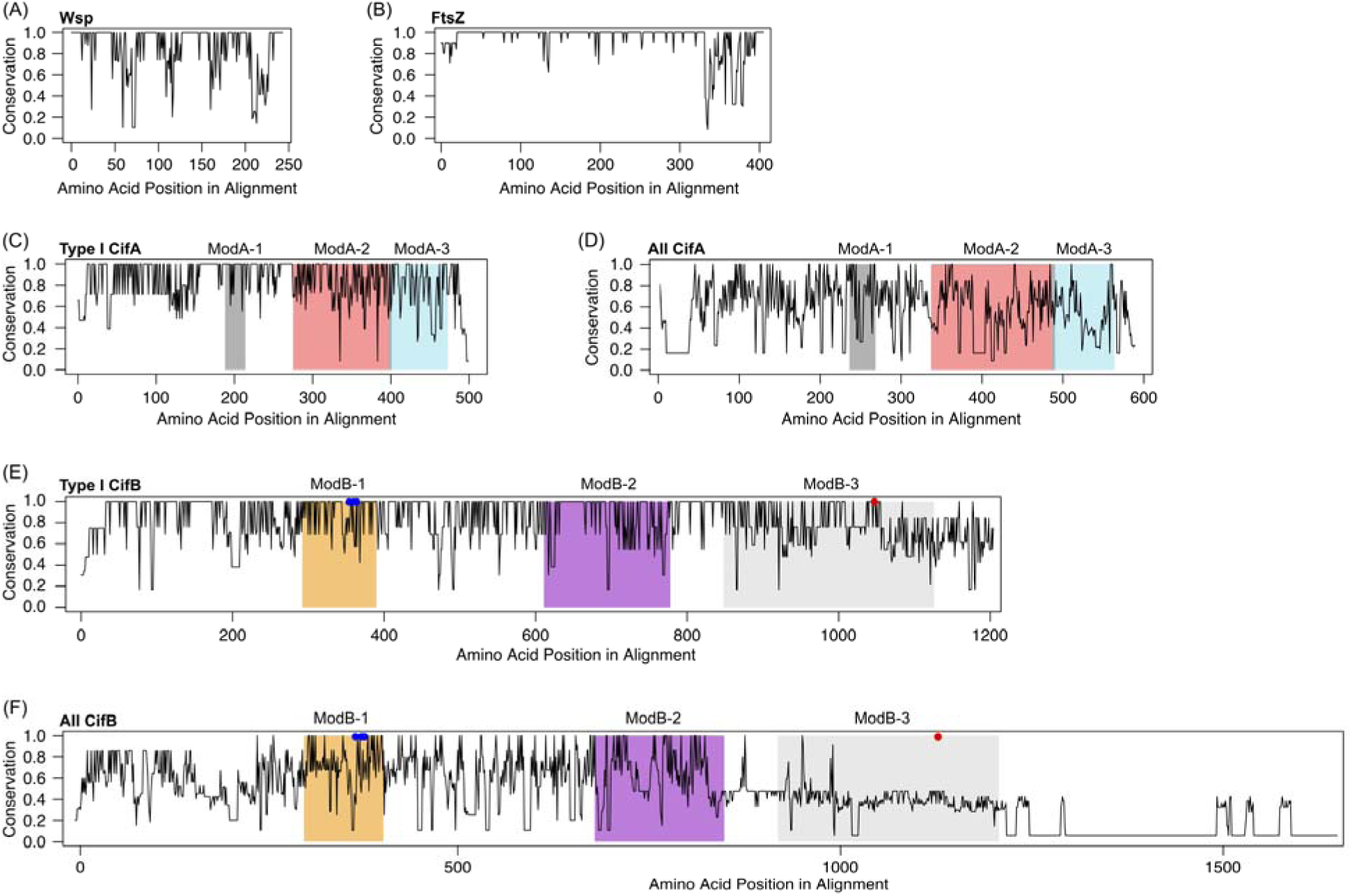
Protein conservation, as determined by Shannon entropy scores. A) Wsp (*Wolbachia* surface protein), B) Cell division protein FtsZ, C) Type I CifA, D) All CifA, E) Type I CifB alleles except for WOVitA4, F) All CifB alleles. Red dots in E and F indicate the ModB-3 catalytic residue (Beckmann, et al. 2017), unique to and completely conserved for Type I alleles. Blue dots in E and F represent the (P)D-(D/E)XK motif (Kosinski, et al. 2005) present in *w*Mel. We found no (P)D-(D/E)XK putative catalytic motif in the second PDDEXK-like module of CifB.

**Table 3.** Average amino acid conservation of Cifs and modules.

| Protein | Region<sup>a</sup> | Type I | All |
| --- | --- | --- | --- |
| CifA | ModA-1 | 0.94 | 0.70 |
|  | ModA-2 | 0.82 | 0.55 |
|  | ModA-3 | 0.77 | 0.53 |
|  | CifA | 0.83 | 0.58 |
| CifB | ModB-1 | 0.89 | 0.71 |
|  | ModB-2 | 0.85 | 0.62 |
|  | ModB-3<sup>b</sup> | 0.77 | 0.39 |
|  | CifB | 0.82 | 0.43 |

<sup>a</sup>Module number defined in Table 2
<sup>b</sup>Only present in Type I

### Cif Module Presence Generally Predicts Reproductive Phenotype

We used the *w*Mel predicted Cif modules as a seed to search for the presence of homologous modules across *Wolbachia* genome sequences using tblastn (Figure 6). In strains with more divergent Cif Types, we report modules that were expected based on the HHpred results, but not recovered with tblastn due to sequence divergence from WOMelB. For example, the WOSuziC and WORiC ModA-1 (Catalase-rel in *w*Mel and other Type I, VirJ in Type II and III) was not recovered. Additionally, we recover homologous modules outside of the annotated *cif* open reading frames, such as the chromosomal region with a ModB-3 (Ulp-1-like) region in *w*No. The high number of modules in *w*Suzi and *w*Ri are due to the presence of a duplicated set of Type I *cifs*. All arthropod-infecting strains, with the exception of *w*Au (a non-CI inducing strain), contained at least one recovered module. This includes the bed-bug mutualist *w*Cle, found in Supergroup-F, and two strains that have lost CI abilities, *w*Uni and *w*Tpre. Importantly, all strains that are known to be capable of inducing or rescuing CI have four or more recovered modules, though they do not necessarily have ModB-3, which contains the catalytic residue implicated in CI function (Beckmann, et al. 2017). The non-CI strains have fewer recovered modules: ModB-1 in *w*Tpre, ModB-1 and −2 in *w*Uni, ModA-3 in *w*Cle. and no modules in *w*Au and the nematode-infecting strains. *w*Uni is a unique case, where we identified *cif* alleles in the genome, but recovered relatively few modules. The CifA modules are either missing (Figure 4A) or divergent enough from WOMelB that they were not considered a positive match. The two N-terminal *w*Uni CifB modules, ModB-1 and ModB-2, are relatively more conserved, and the ModB-3 is missing due to the truncated C-terminus present in all non-Type I CifB alleles (Figure 4B). wAlbB and *w*No, both CI-inducing strains with Type III and IV alleles, have fewer recovered modules, but this is congruent with the more divergent nature of those Cif types. We recovered many modules in *w*Suzi, which is a strain not known to induce CI (Cattel, et al. 2016; Hamm, et al. 2014). This discrepancy between *cif* presence and absence of a reproductive phenotype might be explained by the disrupted Type II *cifA* in *w*Suzi. The split WOSuziC sequenced was concatenated to allow for a more robust phylogenetic reconstruction (Figure 4), but it is in fact disrupted by a transposase (Conner, et al. 2017). However, having a functional set of Type I *cif* alleles appears to be sufficient for CI-induction in other strains (Beckmann, et al. 2017; LePage, et al. 2017), so it is not clear how inactivation of the Type II alleles here may affect the final CI phenotype. Strain *w*Di, infecting the Asian citrus psyllid *Diaphorina citri*, has no identified reproductive phenotype, but only contains a single module: ModB-1.

**Fig 6.**
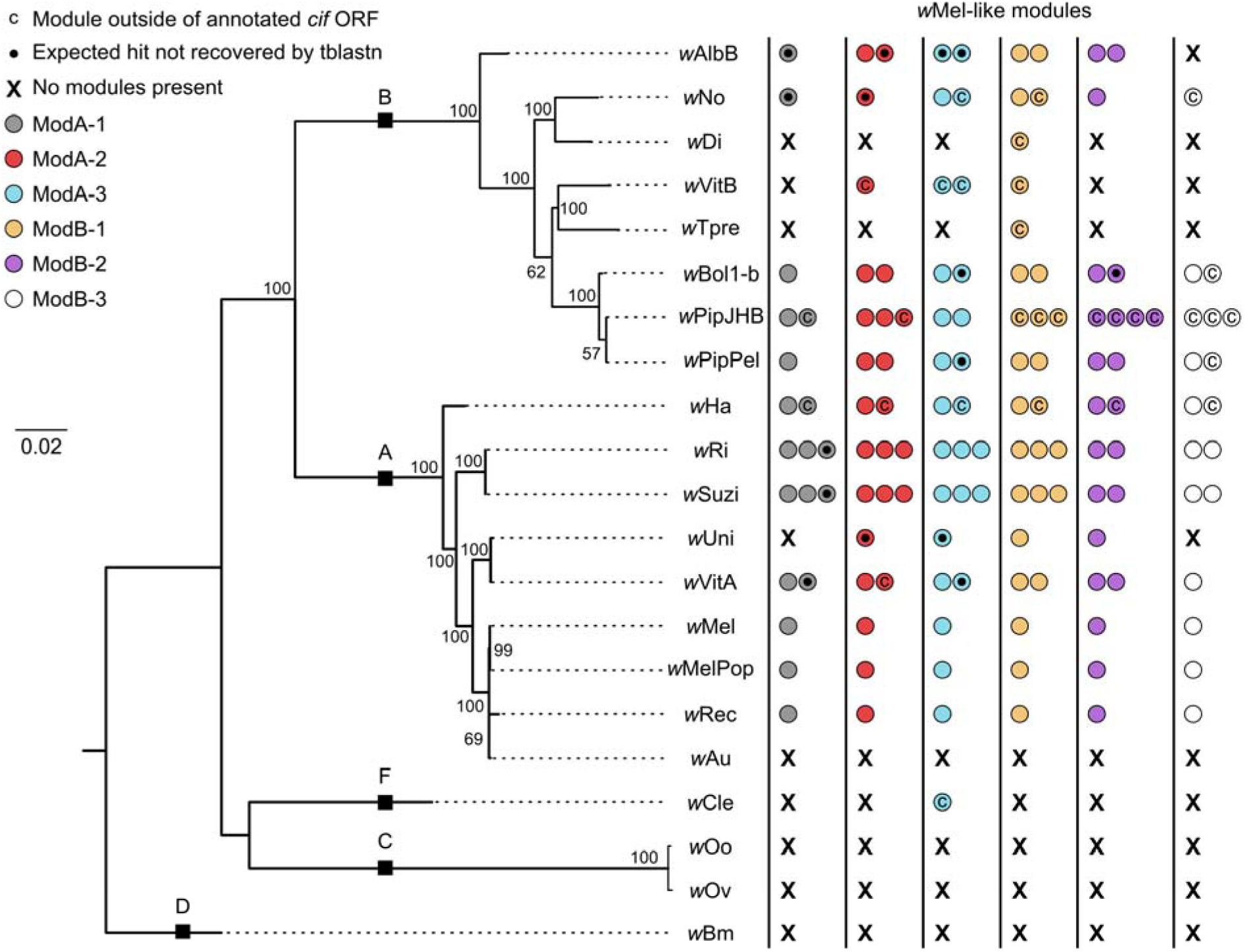
Presence of *w* Mel-like Cif modules across the *Wolbachia* phylogeny. The WOMelB module sequences were used to query available *Wolbachia* genomes to look for the presence of Cif-like regions beyond those within the annotated Cifs (Figure 4). Colored dots correspond to the structural regions delimited by HHpred, shown in Figure 4, and listed in Table 2. A "C" within a dot indicates the presence of a module outside of annotated *cif* open reading frames (Figure 4 and Supplemental Figures S2 and S3). The black dot indicates a module annotated by HHpred, but not identified by tblastn due to divergence from the WOMelB module. Black boxes labeled with uppercase letters indicate branches leading to *Wolbachia* Supergroups. Dotted lines on the phylogeny lead to taxon names and are not included in the branch length.

The lack of evidence for homologous *cif* genes in the nematode-infecting *Wolbachia* agrees with previous findings (LePage, et al. 2017) that CI-function is restricted to the A+B-Supergroup clade (likely due to WO phage activity), and the absence of WO phages for the nematode-infecting strains (Gavotte, et al. 2007). The loss of CI within the A and B Supergroups is likely a derived trait due to the rapid evolution of prophage WO (Ishmael, et al. 2009; Kent, et al. 2011b), and relaxed selection after transition to a new reproductive phenotype. The low number of modules identified in such strains is consistent with gene degradation and loss.

To further explore the conservation of the *cif* genes across the sequenced *Wolbachia,* and to uncover diversity that may be present in other genomes, we searched the WGS databases for recently sequenced genomic scaffolds from *Wolbachia* infecting the *Nomada* bees (*w*Nleu, *w*Nla, *w*Npa, *w*Nfe) (Gerth and Bleidorn 2016), *Drosophila inocompta* (*w*Inc_Cu)(Wallau, et al. 2016), and *Laodelphax striatellus* (*w*Stri) (GenBank Accession Number NZ_LRUH00000000.1) using HMMER. Only for *w*Stri do we have direct evidence of CI induction (Noda, et al. 2001) yet the *w*Stri and *w*Inc_Cu WGS projects each contain only one *cif* locus, with distant homology to *cifA* (∽25% identify across 60% of the *w*Mel protein). Based on HHpred analyses, the *w*Stri homolog (WP_063631193.1) contains none of the domain modules associated with *cifA.* The *w*Inc_Cu homolog (WP_070356873.1) contains three modules: an N-terminal Catalase-rel domain and an internal Ectatomin domain, followed by the STE like transcriptional factor domain. Because these are incomplete genome projects, it is possible that other *cif* homologs have been missed due to the current sequencing coverage. Alternatively, it is possible that other, as yet undiscovered, mechanisms of reproductive manipulation exist in these strains. In contrast, the *Nomada-*associated *Wolbachia* contain a large repertoire of *cif* homologs, including Type I, II, III, IV and several homologs with variations on the Type IV domain architecture for *cifA* (Supplemental Figure S4). The *Nomada Wolbachia* all harbor Type II *cifB* homologs and each of the strains harbors either duplicates of this *cifB* type or novel domain architectures for *cifB* including an N-terminal Oleosin domain and a C-terminal Ulp-1 domain (Supplemental Figure S4).

## Discussion

We explored three key features of *cif* evolution: (i) the toxin-antitoxin operon hypothesis, (ii) potential enzymatic and regulatory functions across the *cifA* and *cifB* phylogenies, and (iii) the conservation and diversity of *cif* genes across strains with different host-manipulation phenotypes. We provide multiple lines of evidence that *cifA* and *cifB* do not comprise an operon in *w*Mel, including quantifications of transcription and *in silico* operon predictors. Moreover, expression of *cifA* and *cifB* across host development are not correlated with each other. In fact, *cifB* expression does not significantly correlate with any other *Wolbachia* locus. Combined with the drastic drop off in expression across the short junction between *cifA* and *cifB*, and negative results from the operon prediction software, we conclude that *cifA* and *cifB* are not co-transcribed or co-regulated as an operon in *w*Mel, the *Wolbachia* strain currently used in mosquito control programs. While we think it unlikely that the *cif* genes are regulated and transcribed in drastically different ways across closely related *cif* Types, more detailed analyses from a variety of strains would be beneficial for developing a comprehensive understanding of the factors regulating expression of these genes. It is especially interesting that synteny has generally been maintained across prophage WO regions, despite the high level of recombination and rearrangements in prophage WO and *Wolbachia* genomes (Baldo, et al. 2006a; Ellegaard, et al. 2013; Kent, et al. 2011a). It is not clear if there is an advantage (and what the advantage may be) to maintaining syntenic orientation of these two genes; perhaps this feature can be attributed to their location within prophage WO and/or functions associated with the ability of *cifA* and *cifB* to act synergistically to induce CI (LePage, et al. 2017). Since type IV secretion system genes and their predicted effectors are scattered across the *Wolbachia* genome (Rice, et al. 2017; Wu, et al. 2004) gene products involved in *Wolbachia-*host interactions can function together even when the genes encoding them are not syntenic. We conclude that *cifA* and *cifB* do not comprise an operon, and do not act strictly as a toxin-antitoxin system due to the requirement of both proteins for the induction of CI in the insect host. Determining how *cifA* and *cifB* expression is regulated in the insect host will greatly benefit vector control programs that use *Wolbachia*-mediated CI.

Despite the conservation of gene order, Cif proteins showed extensive amounts of divergence and differences in domain structure as previously reported (LePage, et al. 2017). Here, the levels of amino acid conservation in the Cifs are lower than FtsZ and Wsp, the latter of which is known to recombine and be subject to directional selection. The conservation of the catalytic residue in the C-terminal deubiquitylase domain is an important feature of CidB (Beckmann, et al. 2017). However, only Type I of the four identified Types has this domain. Additionally, strains known to induce CI, such as *w*AlbB and *w*No have no Type I alleles, implying that the Ulp-1 region may not be essential for inducing CI. The complete, functional capacity of Types I-IV has yet to be explored *in vivo*, but is a promising direction for understanding the evolution of *Wolbachia*-host associations.

Based on what is known about *Wolbachia* biology, some of the protein domains may be especially good candidates for further study and *in vivo* functional characterization. Predicted PDDEXK-like domains are present in all four CifB types. Given the predicted interaction of these domains with DNA (Kosinski, et al. 2005), and the presence of these domains across CifB proteins, determining if and how these regions interact with host (*Wolbachia* or insect) DNA, and whether or not they contribute to CI function would be useful in understanding the consistent presence of this module. Another good candidate for further exploration is the predicted methyltransferase domain in several Type III CifB proteins, as *Wolbachia* infection has been linked to changes in host genome methylation in several insects (LePage, et al. 2014; Negri, et al. 2009; Ye, et al. 2013), though knockout of *Drosophila* methyltransferases does not alter CI levels (LePage, et al. 2014). Likewise, the antioxidant catalase domain is noteworthy as these domains decompose hydrogen peroxide into water and oxygen and thus protect cells from its toxic effects, which are present in *Wolbachia*-infected spermatocytes (Brennan, et al. 2012).

*Wolbachia* strains that have lost CI have a strong signature of *cif* gene degradation and loss. The two parthenogenesis-inducing strains (*w*Tpre and *w*Uni) appear to be at different places in this process of gene loss, with divergent Cif amino acid sequences recovered for *w*Uni, but only one PDDEXK module identified in *w*Tpre. There are several explanations for this. *w*Uni is likely a more recent transition to parthenogenesis, as it is closely related to a CI strain (*w*VitA) (Baldo, et al. 2006b; Newton, et al. 2016). In comparison, *w*Tpre is part of a unique clade of *Wolbachia* that all induce parthenogenesis in *Trichogramma* wasps (Rousset, et al. 1992; Schilthuizen and Stouthamer 1997; Werren, et al. 1995). This strain has lost its WO phage association and only has relics of WO phage genes (Gavotte, et al. 2007; Lindsey, et al. 2016). Additionally, the two strains that independently transitioned to the parthenogenesis phenotype have evolved separate mechanisms for doing so (Gottlieb, et al. 2002; Stouthamer and Kazmer 1994). Differences in time since transition to the parthenogenesis phenotype, phage WO associations, and mechanisms of parthenogenesis induction likely all play a role in the rate of *cif* gene degradation.

Based on our analyses, we propose three avenues of research on the function of the Cif proteins. First, functional confirmation of the newly annotated modules will be important to understanding how these genes function enzymatically. In total, we predict six modules in the Cif protein sequence homologs, with varying degrees of confidence (Supplemental Tables S1 and S2 Tables). For some of these modules, straightforward experiments can be designed in model systems (such as *Saccharomyces*) to determine if their predicted function is correct, as has been done for CidB (Beckmann, et al. 2017) and countless other bacterial effectors (Archuleta, et al. 2011; Kramer, et al. 2007; Siggers and Lesser 2008). Second, necessity and importance of these modules to the CI phenotype can be assessed in the *Drosophila* model, where the induction of the phenotype and rescue is straightforward (LePage, et al. 2017). Finally, we suggest that although the discovery of these genes is fundamental, it is clear from this analysis that we have not comprehensively evaluated or identified the mechanisms behind CI and other reproductive manipulations. The gene characterization analyses described here reveal new and relevant annotations, substantial unknown sequence regions across all of the phylogenetic types, missing deubiquitylase domains in particular CI strains, and a coevolving, phylogenetic relationship across the Cif trees. Importantly, the locus and mechanism behind rescuing CI are still unknown, as is the exact mechanism by which all Cif proteins induce CI. Therefore, the recent discovery of these genes, and the gene characterization analyses described here, pave the most comprehensive road to date for investigating key mechanisms of the *Wolbachia*-host symbiosis.

## Acknowledgments

This work was supported by the National Science Foundation (DEB 1501227 to A.R.I.L., IOS 1456545 to I.L.G.N., and IOS 1456778 to S.R.B); the United States Department of Agriculture (NIFA 2016-67011-24778 to A.R.I.L.); the National Institutes of Health (R21 HD086833 and R01 AI132581 to S.R.B.); and Robert and Peggy van den Bosch Memorial Scholarships to A.R.I.L. We thank J. Dylan Shropshire for feedback on an earlier draft of the manuscript.

